# Characterization of resistance to a potent D-peptide HIV entry inhibitor

**DOI:** 10.1101/748111

**Authors:** Amanda R. Smith, Matthew T. Weinstock, Amanda E. Siglin, Frank G. Whitby, J. Nicholas Francis, Christopher P. Hill, Debra M. Eckert, Michael J. Root, Michael S. Kay

**Affiliations:** Department of Biochemistry, University of Utah School of Medicine, Salt Lake City, UT 84112 USA; Department of Biochemistry and Molecular Biology, Sidney Kimmel Medical College, Thomas Jefferson University, Philadelphia, PA 19107 USA

**Author notes:** Electronic addresses. These authors contributed equally to this work (Amanda R. Smith, Matthew T. Weinstock).

**Keywords:** HIV entry inhibition, D-peptide, PIE12-trimer, gp120, gp41, viral resistance mutation, hydrophobic pocket, viral fitness, compensatory mutation, viral passaging

## Abstract

**Background:** PIE12-trimer is a highly potent D-peptide HIV-1 entry inhibitor that broadly targets group M isolates. It specifically binds the three identical conserved hydrophobic pockets at the base of the gp41 N-trimer with sub-femtomolar affinity. This extremely high affinity for the transiently exposed gp41 trimer provides a reserve of binding energy (resistance capacitor) to combat resistance via stepwise accumulation of modest affinity-disrupting mutations. Viral passaging in the presence of escalating PIE12-trimer concentrations ultimately selected for PIE12-trimer resistant populations, but required an extremely extended timeframe (>1 year) in comparison to other entry inhibitors. Eventually, HIV developed resistance to PIE12-trimer by mutating Q577 in the gp41 pocket.

**Results:** Using deep sequence analysis, we identified three mutations at Q577 (R, N and K) in our two PIE12-trimer resistant pools. Each point mutant is capable of conferring the majority of PIE12-trimer resistance seen in the polyclonal pools. Surface plasmon resonance studies demonstrated substantial affinity loss between PIE12-trimer and the Q577R-mutated gp41 pocket. A high-resolution x-ray crystal structure of PIE12 bound to the Q577R pocket revealed the loss of two hydrogen bonds, the repositioning of neighboring residues, and a small decrease in buried surface area. The Q577 mutations in an NL4-3 backbone decreased viral growth rates. Fitness was ultimately rescued in resistant viral pools by a suite of compensatory mutations in gp120 and gp41, of which we identified seven candidates from our sequencing data.

**Conclusions:** These data show that PIE12-trimer exhibits a high barrier to resistance, as extended passaging was required to develop resistant virus with normal growth rates. The primary resistance mutation, Q577R/N/K, found in the conserved gp41 pocket, substantially decreases inhibitor affinity but also damages viral fitness. Identified candidate compensatory mutations in gp160 will be the focus of future mechanistic studies.

## Background

Approximately 37 million people are living with Human Immunodeficiency Virus (HIV), a virus that has resulted in an estimated 35 million deaths due to Acquired Immunodeficiency Syndrome (AIDS) [1]. Combination antiviral therapeutics targeting various stages of the viral lifecycle have dramatically improved the clinical outcome for those with access to treatment, but viral resistance has been documented for all approved drugs [2]. This resistance is due to a low-fidelity viral reverse transcriptase, frequent recombination events, rapid turnover of the virus, and host APOBEC3 activity[3–5].

We previously reported the discovery and design of an extremely potent D-peptide HIV-1 entry inhibitor (PIE12) that targets the HIV envelope protein (Env) using sequential rounds of mirror-image phage display and high-resolution structure-guided design [6–9]. With the opposite chirality of naturally occurring L-peptides, D-peptides (composed of D-amino acids) are not recognized by natural proteases. This property confers longer in vivo half-lives and reduced immunogenicity, which are potentially significant therapeutic advantages [7]. Our inhibitor harnesses avidity for the highly conserved trimeric pocket at the base of the HIV-1 gp41 N-trimer, as three D-peptide monomers are covalently linked via an optimized PEG-based scaffold (PIE12-trimer), and binds the N-trimer with sub-fM affinity [8, 10]. PIE12-trimer prevents the collapse of the N-trimer and C-peptide regions of gp41 into the 6-helix bundle (trimer-of-hairpins) structure required for membrane fusion during viral entry, and it inhibits all members of a diverse panel of primary Group M HIV isolate strains with high pM to low nM potency (IC50) [8].

The remarkable conservation of this gp41 hydrophobic pocket target [6] is indicative of its critical roles in the viral life cycle and impedes the ability of the virus to combat inhibitors that target it. The region of the HIV-1 genome encoding the pocket is conserved at both the protein level (for the role of gp41 in membrane fusion during viral entry) and nucleotide level (due to a structured RNA region in the critical Rev-response element). The only FDA-approved HIV fusion inhibitor, enfuvirtide (Fuzeon), is an L-peptide derived from the gp41 C-peptide region that targets the gp41 N-trimer region adjacent to, but not including, the hydrophobic pocket and induces the rapid emergence of gp41 resistance mutations both in vitro and in vivo [11, 12]. Next-generation L-peptide fusion inhibitors (such as T-1249 and T-2635), whose binding footprint extends into the pocket, are much less impacted by enfuvirtide resistance mutations. Viral escape from these inhibitors generally takes longer to appear but still involves resistance mutations located outside the pocket region [13–16].

In addition to targeting a highly conserved, functionally critical region of the virus, PIE12-trimer was designed with an additional barrier to viral resistance called the “resistance capacitor”. The gp41 N-trimer is transiently exposed during HIV entry in a prehairpin intermediate conformation [17]. The potency of a variety of high-affinity prehairpin intermediate-targeting inhibitors has been shown to vary with changes in association kinetics rather than affinity, reflective of the kinetically limited availability of the target [8, 9, 18–20]. Once a threshold association rate is reached, additional affinity enhancement fails to improve antiviral potency. We chose to “over-engineer” PIE12-trimer such that its pocket-binding affinity far surpassed its potency. This excess binding energy, or resistance capacitor, should prevent the accumulation of incremental mutations that modestly disrupt inhibitor affinity, as such mutations would not affect potency and would therefore fail to confer a selective replicative advantage. Instead, the virus would be forced to combat PIE12-trimer with a strongly disruptive mutation, likely at the cost of viral fitness.

Several lines of evidence suggested that PIE12-trimer would exhibit a high barrier to resistance. First, PIE12-trimer is estimated to bind with sub-fM affinity and inhibit with low nM potency, characteristic of a “highly charged” resistance capacitor [8]. Second, PIE12-trimer exhibits higher potency against polyclonal viruses derived from infected patients than does an earlier generation D-peptide trimer with lower affinity, demonstrating its better tolerance of Env sequence variation. Finally, PIE12-trimer maintains potency against virus resistant to PIE7-dimer, a lower affinity D-peptide. Indeed, consistent with a “highly charged” resistance capacitor, we observed a significant delay in the selection of HIV-1 resistant to PIE12-trimer compared to selections against enfuvirtide and earlier-generation D-peptides [8].

Expanding on our earlier work that selected for viral resistance to PIE12-trimer, here we describe the biochemical, structural, and functional properties of the primary PIE12-trimer resistance mutations (Q577R/N/K). These mutations substantially disrupt PIE12-trimer affinity and significantly decrease viral fitness. Through comprehensive characterization of the genotype of our PIE12-trimer resistant viral populations via next-generation sequencing, we identify potential compensatory mutations in HIV Env that restore fitness in the context of Q577R/N/K.

## Methods

### Passaging Studies

PIE12-trimer resistant viruses were acquired in a previous study [8]. Briefly, CEM-1 cells were infected with the laboratory-adapted HIV-1 strain NL4-3 (obtained from Dr. Malcolm Martin, Cat #114, NIH AIDS Reagent Program). Virus was serially propagated weekly by adding a 1:5 dilution of cell-free viral supernatant to fresh CEM-1 cells in the presence or absence (control) of PIE12-trimer. Viral titers were monitored biweekly by p24 antigen ELISA (PerkinElmer). The initial inhibitor concentration was 1 nM (near the PIE12-trimer IC50). When p24 antigen levels in PIE12-trimer-containing cultures approached the level seen in inhibitor-free (control) cultures (typically 5-15 weeks), PIE12-trimer concentrations were raised approximately 1.5-to 2-fold. Ultimately, two polyclonal viral pools able to propagate in the presence of 320 nM PIE12-trimer (W1 and W2) were obtained. These pools were propagated in CEM-GFP cells for 48 h then isolated and concentrated over a 20% sucrose cushion.

### Cells and Viruses

TZM-bl cells that express luciferase and β-galactosidase in response to HIV-1 promoter activation (obtained from Dr. John C. Kappes and Dr. Xiaoyun Wu, Cat #8129, NIH AIDS Reagent Program) and 293T cells were grown in Dulbecco’s modified Eagle media. CEM-GFP cells that express GFP in response to HIV-1 LTR activation (obtained from Dr. Jacques Corbeil, Cat #3655, NIH AIDS Reagent Program) were grown in RPMI 1640 media. The Q577R, Q577N and Q577K point mutant viral vectors were generated by excising the *env* region of the WT pNL4-3 genome using XhoI and EcoRI (NEB) and ligating into pET20b (Novagen), a shuttle vector we used for mutagenesis. The point mutants (CGG for Q577R, AAG for Q577K and AAC for Q577N) were introduced using the Q5 site-directed mutagenesis kit with primers designed using NEBuilder software (NEB). The Q577R forward and reverse primers were CAAACAGCTCCGGGCAAGAATCC and ATGCCCCAGACTGTGAGTT. The Q577N forward and reverse primers were CAAACAGCTCAACGCAAGAATCCTGG and ATGCCCCAGACTGTGAGT. The Q577K forward primer was CAAACAGCTCAAGGCAAGAATCC, and the reverse primer was the same as that for Q577N. Mutated clones were confirmed by Sanger sequencing, and the desired mutants were ligated back into the WT pNL4-3 backbone after EcoRI/XhoI digestion. *Env* in the final Q577-mutated pNL4-3 constructs was confirmed by Sanger sequencing. These vectors were then transfected into 293T cells using Fugene HD (Promega) to generate replication-competent virus. Viral titers were measured using the HIV-1 p24 Antigen Capture Assay (Advanced Bioscience Laboratories) and confirmed by X-gal staining of infected TZM-bl cells.

### Generation of envelope cDNA for Sanger and deep sequencing

Viral RNA was isolated from the W1, W2, and control viral pools with the E.Z.N.A.® Viral RNA Kit (Omega Bio-Tek). *Env* cDNA was generated using the SuperScript® III One-Step RT-PCR System with Platinum® *Taq* High Fidelity (Invitrogen). Barcoded primers were designed using Primer3Plus (RT Primer: GCCTTGCCAGCACGCTCAGGCCTTGCATAGTAGCNNNNNNNNATGAGAGTGAAGGAGAAGTATCAGC and PCR primer FWD: GCCTTGCCAGCACGCTCAGGC) [21]. For Sanger sequencing, the *env* cDNA was cloned into standard vectors via two strategies. First pEBB backbone was used with NotI/XhoI sites. For increased efficiency, we also used TOPO cloning. Individual plasmid clones were sequenced by the DNA Sequencing Core Facility (University of Utah).

For deep sequencing, the cDNA was purified by agarose gel electrophoresis, extracted (GenElute™ Gel Extraction Kit, Sigma), and submitted to the High-Throughput Genomics and Bioinformatic Analysis Core Facility (Huntsman Cancer Institute, University of Utah) for library preparation and sequencing. Barcoded samples were multiplexed in a single lane and sequenced on an Illumina HiSeq 2000 (50 cycle single-end reads). The raw sequencing reads in FASTQ format were mapped to nucleotides 6195-8826 from the HIV-1 vector pNL4-3 sequence (GenBank: AF324493.2) using STAMPY [22]. A summary of base calls for each position of the reference sequence was generated by converting the alignment data to pileup format using the SAMtools:*mpileup* function (used -BQ0 and -d 2000000 options). Custom Perl scripts were used to convert base calls at each position from pileup format to a more intuitive table and then to filter the data to display only those positions where deviations from the reference are present in the population above a user-defined threshold (10%).

### Peptide Synthesis

All synthetic peptides were acetylated at the N-terminus and amidated at the C-terminus. All reverse-phase HPLC (RP-HPLC) purifications were performed on a Waters C18 column (BEH X-Bridge, 19 x 250 mm, 10 µm, 300 Å) with water/acetonitrile gradient in 0.1% TFA unless otherwise noted. KG-PIE12 was synthesized by the University of Utah DNA/Peptide Core as described in [8] and purified via RP-HPLC on a Vydac C18 column (TP, 22 x 250 mm, 10 µm, 300 Å). PIE12-GK and PIE12-trimer were synthesized as previously described [8]. SelMet-IQN17-Q577R and HIV C34 were synthesized on a PS3 peptide synthesizer (Protein Technologies, Inc.) at 80 µmol scale on NovaPEG Rink Amide LL resin (Novabiochem) and Rink Amide AM resin LL (Novabiochem), respectively, via Fmoc-SPPS and were purified by RP-HPLC.

### Recombinant protein production

Cys-Gly-Gly-Asp-IZN36 (derived from HXB2 strain) with a thrombin-cleavable His tag was prepared as previously described [8]. A Q577R mutant version was generated via site-directed mutagenesis using the QuikChange® protocol (Stratagene). Protein expression was performed in BL21 (DE3) cells (Novagen) following Studier’s autoinduction protocol [23]. Inclusion bodies containing IZN36 were solubilized in 20 mM sodium phosphate pH 8.0, 300 mM NaCl, 10 mM imidazole, 6 M GuHCl, and sonicated. IZN36 was then purified using His-Select Ni-affinity resin (Sigma). Ni-purified samples were dialyzed into water with 0.1% TFA and purified via RP-HPLC on a Waters C18 column (BEH X-Bridge, 19 x 250 mm, 10 µm, 300 Å) with water/acetonitrile gradient in 0.1% TFA and lyophilized. The cysteine residues of these constructs were biotinylated for SPR. Reactions were carried out in PBS pH 7.0 with ∼1-2 mg/mL protein and a 15X molar excess of Biotin-dPEG®3-MAL (Quanta BioDesign) at room temp for 1 hr, followed by RP-HPLC purification. The His tag was cleaved in PBS pH 6.3 with 10 µM heparin at a protein concentration of ∼0.8 mg/mL and one unit of thrombin protease (GE Healthcare) per mg of protein. The cleavage reaction was incubated at 37°C with agitation for 44 hrs followed by RP-HPLC as above.

### Surface Plasmon Resonance Analysis

Protein interaction analysis was carried out on a Biacore 3000 (GE Healthcare). Analyses were performed in triplicate at 25°C in sterile-filtered and degassed Gibco® PBS pH 7.4 (Invitrogen) supplemented with 0.005% P20 and 1 mg/mL BSA at a flow rate of 50 μL/min. First, streptavidin (Pierce) was immobilized on CM5 chips (GE Healthcare) using the Amine Coupling Kit (GE Healthcare) following a standard coupling protocol [24]. Stocks (∼50 nM) of biotinylated IZN36 (WT and Q577R mutant) were prepared in running buffer in the presence of 1.2x molar excess of HIV C34 and captured on the surface to a density of 100-200 RUs. Loading IZN36 in the presence of C34 results in better behaved surfaces, and C34 was removed by surface regeneration with 0.1% SDS injections. The active surface and a streptavidin-derivatized control surface were blocked with D-biotin. Stock solutions of PIE12-GK were prepared at 50 μM in running buffer, from which samples in the concentration series were generated by serial dilution (three-fold to 2.54 nM). Injections were performed for 1 min using KINJECT. Dissociation was monitored for 2 min, followed by a 20 μL injection (QUICKINJECT) of running buffer followed by EXTRACLEAN. Sensorgrams were processed using Scrubber2 (BioLogic Software), and data were analyzed with Prism 5.0 (GraphPad Software). The equilibrium binding data were fit using the built-in saturation binding model (one site -- specific binding), normalized to Bmax, and replotted as concentration of analyte vs. % capacity.

### Crystallography

KG-PIE12 was crystallized in complex with Se-Met-IQN17-Q577R (1:1.1 IQN17:KG-PIE12, 10 mg/mL total in water) at 21°C in Hampton Research condition SaltRx HT B4 (1.8 M ammonium citrate dibasic, 0.1 M sodium acetate trihydrate, pH 4.6) by sitting-drop vapor diffusion. Crystals were mounted in a nylon loop and either directly cryocooled by plunging into liquid nitrogen or cryocooled following a 20 s immersion in 20 μL crystallization buffer with 20% added glycerol. Crystals were maintained at 100 K during data collection. Data collection was performed at the Stanford Synchrotron Radiation Lightsource (SSRL) beam line 7-1. Data were processed using HKL2000 [25]. The IQN17-Q577R/PIE12 structure was determined by molecular replacement using PHASER [26] with PIE12-IQN17 (PDB structure 3L37) as the search model. The model was rebuilt using O [27] and COOT [28] and refined against a maximum-likelihood target function using PHENIX [29]. The structure was checked using the MolProbity program [30]. See Table 2 for crystallographic data and refinement statistics. Buried surface area was calculated with the program Naccess V2.1.1 [31]. For compatibility with Naccess, the surface areas were calculated with the ACE and NH2 modified ends removed, and the names of the D-amino acid residues changed to their cognate L-amino acids, without changing the coordinates.

### Growth Assay

CEM-GFP cells were infected with 1 ng p24 of either clonal pNL4-3, one of the Q577 point mutant viruses, or one of the polyclonal viral pools (W1 or W2) with DEAE dextran (8 µg/mL) and incubated at 37°C for 2 h. Cells were then pelleted (2000 *x g*, for 5 min), supernatant was removed, and cells were resuspended in fresh media. Cells were incubated for 3 days, then the medium was changed every day for the next 7 days. Viral supernatant was collected on days 3, 5, 7, and 9 post-infection, and RNA was extracted using E.Z.N.A.® Viral RNA Kit (Omega Bio-Tek). Viral levels were measured by qRT-PCR (Lightcycler 480, Roche) with primers targeting a conserved region of Gag and compared to a standard curve of samples with known quantities of p24.

### Viral Inhibition Assays

Viral infectivity was measured as previously described [32], with the following changes. Replication-competent viruses were added to TZM-bl cells in the presence of a 10-fold dilution series of PIE12-trimer (100 nM to 10 pM for WT virus, 100 µM to 10 nM for resistant virus, from PIE12-trimer stock in 50 mM HEPES pH 7.4) and 8 µg/mL DEAE dextran. Medium was removed after 30 hours, and cells were lysed with Glo Lysis buffer (Promega). BrightGlo luciferase substrate (Promega) was added and luminescence was measured (POLARstar Optima, BMG). TZM-bl cells exhibit significant background luminescence, so uninfected wells were used for background subtraction. Luciferase activity was normalized to uninhibited controls, and IC_50_ values were determined from a fit to a Langmuir equation: y=1/(1+[inhibitor]/IC_50_) weighted by the normalized standard error of replicates within an assay (with a minimum 1% error) using Prism 7.04 (GraphPad).

## Results

### Primary Resistance to PIE12-Trimer Arises Via Mutation of Q577 in the gp41 Hydrophobic Pocket

We previously reported the selection of HIV-1 populations resistant to PIE12-trimer by passaging the lab-adapted NL4-3 strain in the presence of escalating suboptimal (partially inhibiting) concentrations of inhibitor, starting with 1 nM. Ultimately, PIE12-trimer resistant virus propagating in 320 nM PIE12-trimer was obtained after 65 weeks of passaging [8]. Duplicate selections yielded two highly resistant polyclonal viral pools (W1 and W2). As previously reported, similar passaging studies with enfuvirtide led to high-level resistance (>1000-fold) within ∼3 weeks [8]. Both W1 and W2 showed >650-fold loss of sensitivity to PIE12-trimer while retaining full susceptibility to the C-peptide inhibitor, C37, demonstrating the ability of the virus to specifically resist PIE12-trimer while maintaining the native interaction between the N-trimer and C-peptide (Fig. 1 and Table 1).

**Figure 1:**
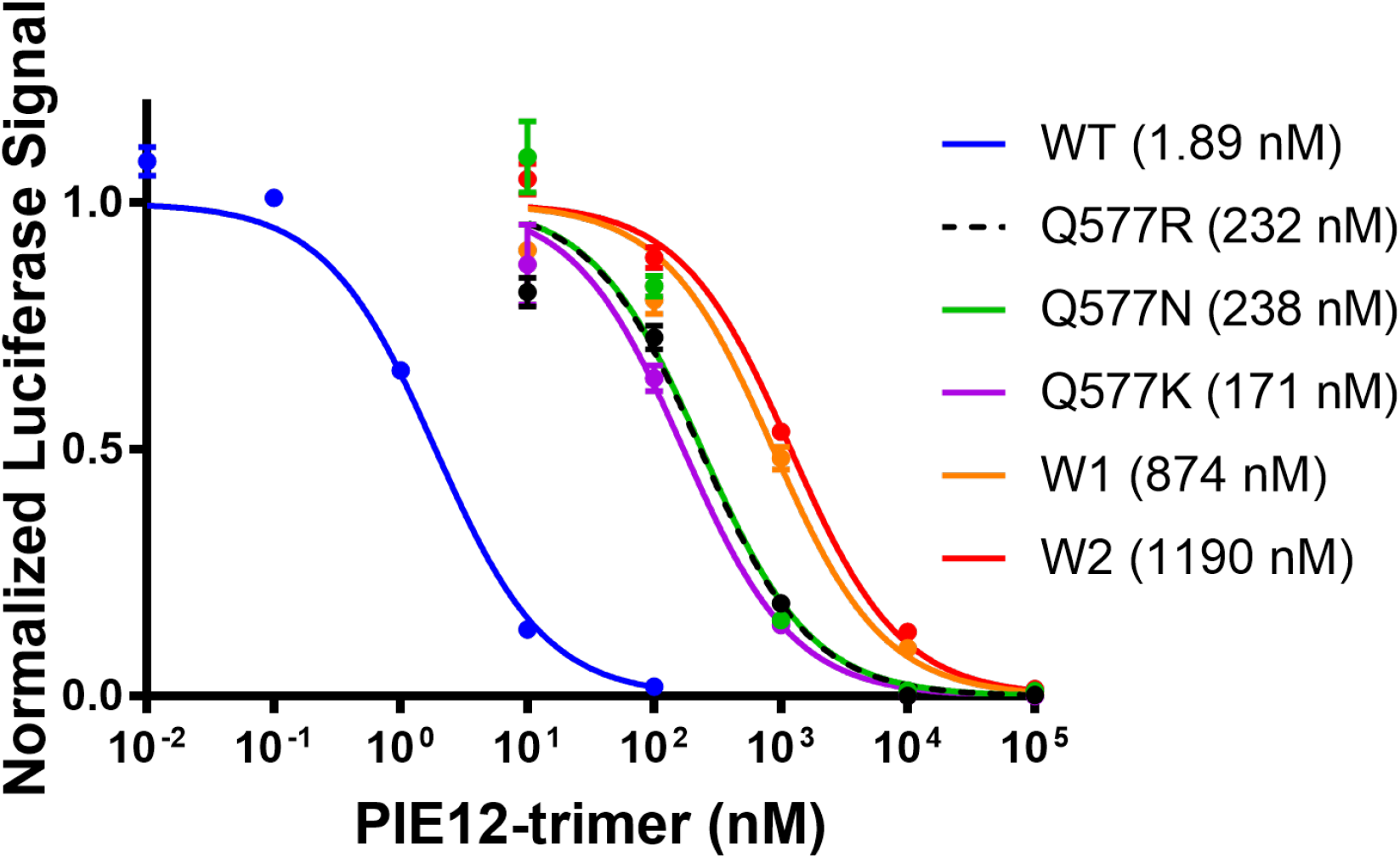
Representative inhibitory curves of replication-competent NL4-3 (WT) and PIE12-trimer resistant viruses. The polyclonal viral pools (W1 and W2) show slightly enhanced resistance compared to clonal virus expressing Env with Q577 point mutations. Representative normalized data are shown from a single assay with triplicate measurements (error bars show standard error). IC_50_ values of this representative data set are indicated in parentheses.

**Table 1:** Inhibition of replication-competent HIV-1 (NL4-3 backbone) Average IC_50_ values for clonal NL4-3 (WT), PIE12-trimer 320 nM polyclonal resistant pools (W1 and W2), and Q577 point mutations in the NL4-3 background. Values are averages of IC_50_’s fit from two independent experiments with triplicate measurements. Standard deviations are shown in parentheses.

|  | PIE12-trimer (nM) | C37 (nM) |
| --- | --- | --- |
| NL4-3 (WT) | 1.33 (0.79) | 11.2 (4.07) |
| W1 (resistant pool) | 999 (176) | 7.62 (2.18) |
| W2 (resistant pool) | 894 (419) | 14.9 (1.31) |
| Q577R | 188 (63) | ND |
| Q577N | 164 (105) | ND |
| Q577K | 128 (59) | ND |

Early analysis of a limited number of clones from the W2 resistant pool via Sanger sequencing identified the mutation Q577R within the gp41 hydrophobic pocket in all clones [8]. In an HXB2 pseudovirion, Q577R confers >4000-fold resistance to PIE12-trimer [10]. Here, to characterize the genetic basis of resistance and to understand the penetrance of the Q577R mutation, we deep sequenced the W1 and W2 viral pools, as well as a pNL4-3 control viral pool that had been propagated in the absence of inhibitors. This more extensive sequencing (with an average sequencing depth of >200,000 reads per nucleotide position) revealed two distinct mutations at position Q577 (codon CAG) in the highly resistant pools: N (codon AAC) in W1 and R (codon CGG) in W2. Q577 is mutated in virtually all of the resistant virus (>97% of the sequences in W1 and W2), and there were no spontaneous Q577 mutations in the control population (Supplementary Table 1). Additionally, there are no other positions within the gp41 pocket that are mutated.

Q577R only requires a single nucleotide change, while Q577N requires a double mutation and likely occurs stepwise. To determine which pathway the virus took to acquire the Q577N substitution, we deep sequenced this N-trimer region from an intermediate pool of virus isolated at an earlier timepoint during resistance selection and noted an equal ratio of N (AAC) and K (AAG) (data not shown). In the context of replication-competent virus (NL4-3 strain), each Q577 mutant (R, N or K) confers substantial resistance to PIE12-trimer (∼100-fold) and accounts for the majority of the loss of sensitivity to PIE12-trimer (Fig. 1 and Table 1).

### Mutation of Q577 Significantly Decreases PIE12-trimer Affinity for the gp41 Pocket

To determine the impact of Q577 mutation on PIE12 binding affinity, Q577R was incorporated into an N-peptide mimic (IZN36 [17]), and the interaction was analyzed via surface plasmon resonance (SPR). Because PIE12-trimer affinity is too tight to accurately measure via SPR[8], monomeric PIE12 was flowed over the IZN36 (WT and Q577R) surfaces. The Q577R mutation results in a dramatic (∼65-fold) weakening of the interaction between PIE12-monomer and the hydrophobic pocket (Q577R K_D_=2.0 µM vs. 0.031 µM for WT) (Fig. 2 and Supplementary Fig. 1), corresponding to a 2.5 kcal/mol weakening of the PIE12 interaction. Because our inhibitor is trimeric, we expect the impact on PIE12-trimer’s binding energy to be three times as great, corresponding to a geometric drop of binding affinity of 65^3^ (or >250,000). This large decrease in binding affinity is consistent with the prediction that an extremely disruptive mutation is required to overcome PIE12-trimer’s highly charged resistance capacitor.

**Figure 2:**
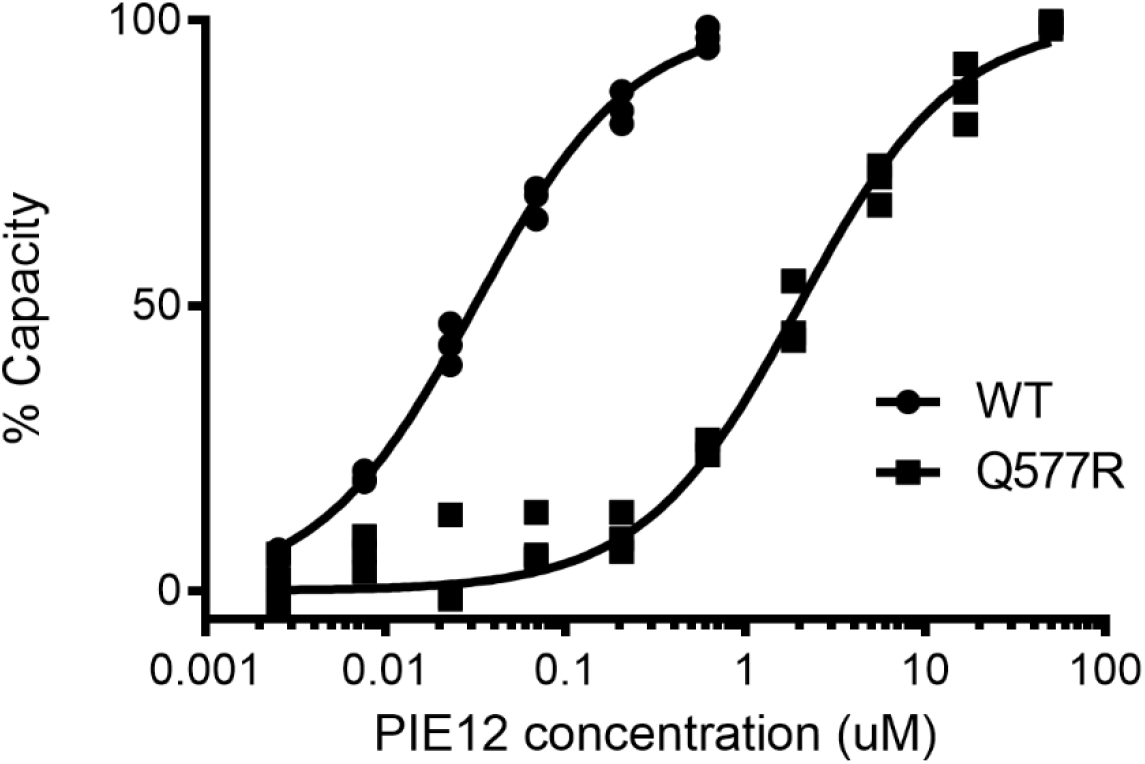
Protein interaction analysis of Q577R mutation. SPR equilibrium binding data (in triplicate) for interaction between immobilized IZN36 (WT or Q577R) and PIE12 monomer. The calculated K_D_ values are 0.031 µM for WT and 2.0 µM for Q577R.

### Crystal Structure of PIE12 in Complex with the Q577R Mutation

To understand the molecular basis of Q577R-induced resistance, monomeric PIE12 was crystallized in complex with an HIV-1 N-trimer pocket mimic (IQN17 [17]) with the Q577R substitution, and the structure was determined at 1.3 Å resolution (Table 2). As in all previous D-peptide/IQN17 structures [6, 8, 9], IQN17 forms a continuous trimeric coiled coil with the D-peptide binding to the gp41 hydrophobic pocket region. When compared to our previously reported 1.45 Å structure of PIE12 binding to WT IQN17 (PDB code 3L37) [8], the overall structure agrees well, with a root mean square deviation (RMSD) of 0.34 Å for all Cα atoms. An overlay of these two structures, centered around position 577 of the hydrophobic pocket is shown in Figure 3. The WT structure shows that Q577 makes hydrogen bond contacts with D-Glu9 and D-Trp12 in PIE12 [8]. Q577R disrupts these hydrogen bonds, likely accounting for the loss of affinity, and induces a repositioning of two adjacent residues (D-Glu9 in PIE12 and L581 in IQN17). These structural changes lead to a minimal change in the buried surface area of the inhibitor interface with the N-trimer target (555 and 524 Å^2^ in the WT and mutant structures, respectively).

**Table 2:** Crystallographic data and refinement statistics.

|  | Q577R |
| --- | --- |
| Data |  |
| Crystal | LA5QR |
| Processing software | HKL2000 |
| Space Group | R3<br>(48.8, 48.8, 67.6) |
| Resolution (Å) | 50.0 – 1.30 |
| Resolution (Å) (high-resolution shell) | (1.32 – 1.30) |
| # Reflections measured | 146783 |
| # Unique reflections | 14574 |
| Redundancy | 10.1 |
| Completeness (%) | 98.5 (99.9) |
| $\langle I/\sigma I \rangle$ | 12 (0.9) |
| Mosaicity (°) | 0.62 |
| Rsym<sup>a</sup> | 0.074 (0.375) |
| Refinement |  |
| Refinement software | PHENIX.REFINE |
| Resolution (Å) | 50.0 – 1.30 |
| Resolution (Å) – (high-resolution shell) | (1.347 – 1.300) |
| # Reflections used for refinement | 14513 |
| # Reflections in Rfree set | 1454 |
| Rcryst<sup>b</sup> | 0.176 (0.312) |
| Rfree<sup>c</sup> | 0.217 (0.305) |
| RMSD: bonds (Å) / angles (°) | 0.008 / 1.123 |
| $\langle B \rangle$ (Å<sup>2</sup>): all protein and peptide atoms / # atoms | 22.7 / 578 |
| $\langle B \rangle$ (Å<sup>2</sup>): water molecules / #water | 40.3 / 74 |
| $\phi/\psi$ most favored (%) / additionally allowed (%) | 97.6 / 2.4 |

Values in parenthesis refer to data in the high-resolution shell.
<sup>a</sup> Rsym = $\sum |I - \langle I \rangle| / \sum I$ where $I$ is the intensity of an individual measurement and $\langle I \rangle$ is the corresponding mean value – precision-indicating merging R-value.
<sup>b</sup> Rcryst = $\sum ||F_o| - |F_c|| / \sum |F_o|$ , where $|F_o|$ is the observed and $|F_c|$ the calculated structure factor amplitude.
<sup>c</sup> Rfree is the same as Rcryst calculated with a randomly selected test set of reflections that were never used in refinement calculations.

**Figure 3:**
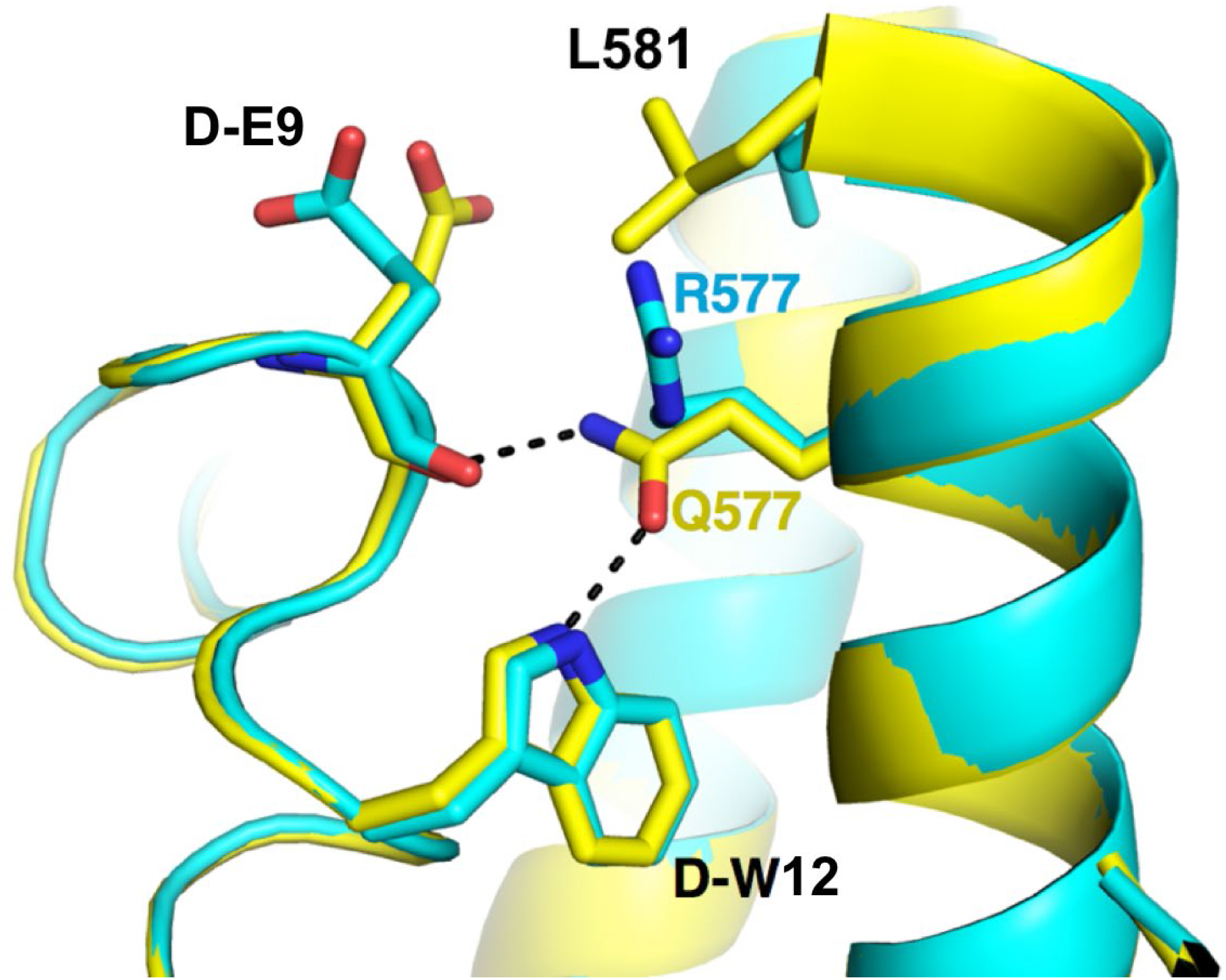
Crystal structures of PIE12/IQN17 (Q577R) and PIE12/IQN17 (WT PDB:3L37). Overlay of crystal structures of PIE12 binding to the WT (yellow) and a mutant (Q577R, turquoise) pocket showing the disruption of hydrogen bonding interactions mediated by Q577.

### Mutation of Q577 Causes a Significant Viral Growth Defect

The extended time required for evolution of high-level PIE12-trimer resistance in passaging studies (65 weeks) and the high sequence conservation of the inhibitor binding site on gp41 suggest that the primary Q577 resistance mutations come with a fitness cost to the virus. We therefore investigated the growth kinetics of the NL4-3-based replication competent Q577 point mutants and the polyclonal viral pools (W1 and W2) and compared them to that of the native virus (clonal NL4-3, WT). Starting with a virus inoculum containing 1 ng p24, indeed, the growth of the Q577 mutants is significantly decreased compared to WT (Fig. 4). Q577R and Q577K did not propagate significantly during the entire course of the assay, while Q577N replication was substantially delayed. Interestingly, the PIE12-trimer resistant polyclonal pools (W1 and W2) show similar growth kinetics to WT virus, suggesting that compensatory mutations rescue the fitness defect associated with the Q577 mutants. WT, W1, and W2 all show rapid growth and hit similar peak p24 levels. Importantly, starting viral titers of the Q577 mutant stocks (normalized to p24) vary significantly from WT, ranging from greatly reduced (∼40-fold for Q577K and ∼7-fold for Q577R) to increased (∼3-fold for Q577N).

**Figure 4.**
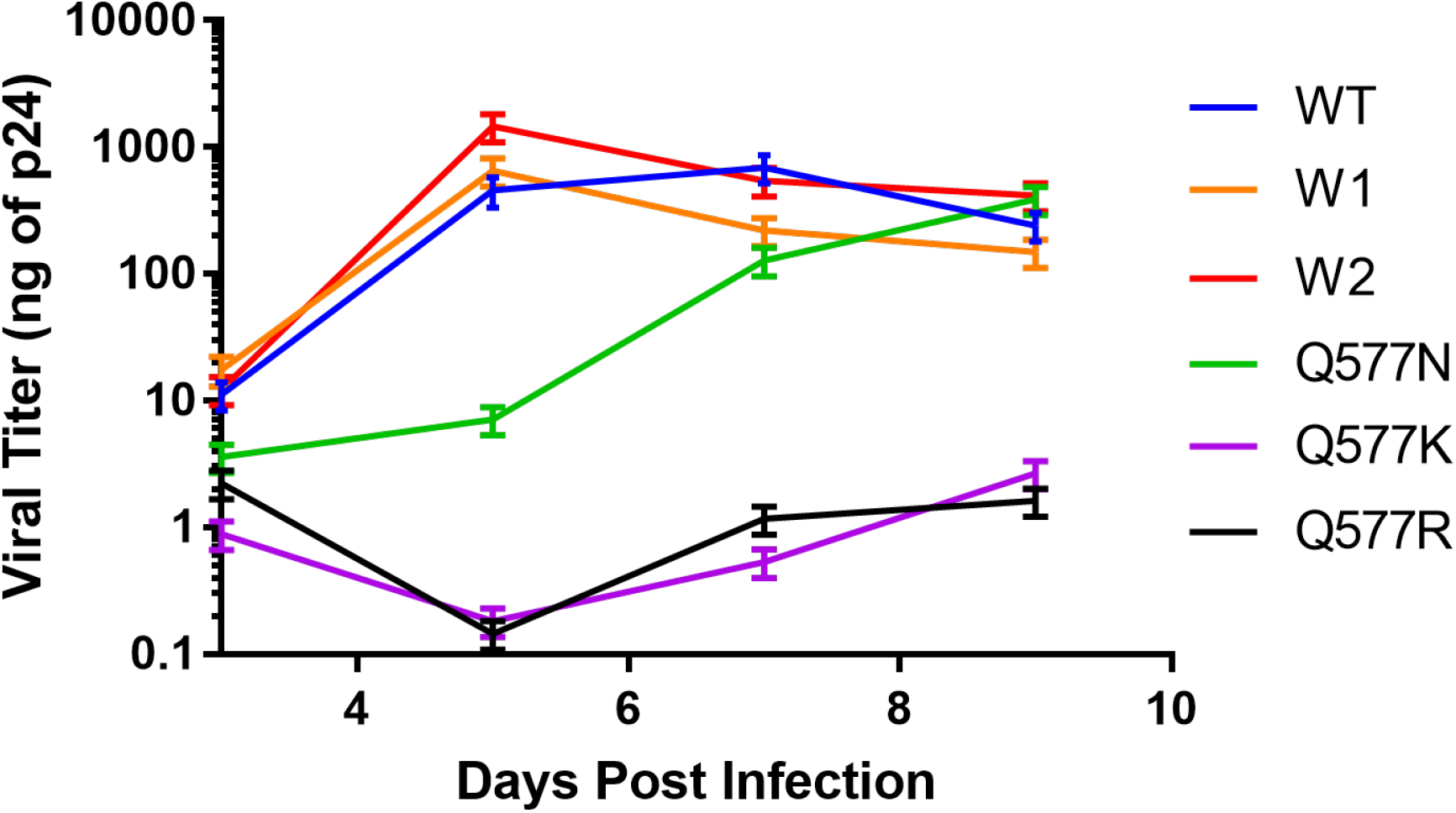
Viral Growth Assays. Comparison of growth kinetics between control clonal NL4-3 (WT), Q577 mutant viruses (R, N and K), and resistant polyclonal viral pools (W1 and W2). Viral titers were determined by qRT-PCR and compared to a standard curve of known HIV-1 titers (based on p24 antigen levels). Each culture was inoculated with virus containing 1 ng p24 (average of biological duplicate assays shown with s.d. error bars).

### Identification of Potential Compensatory Mutations to Overcome the Q577 Fitness Defect

To identify additional mutations that arose during the selection for resistance, the entire *env* gene for each resistant pool (and control pool propagated in the absence of inhibitor) was deep sequenced. To complement these short reads and obtain linkage information, we also performed Sanger sequencing on 13 PIE12-trimer resistant clones (five from W1 and eight from W2). This search should identify mutations that compensate for the fitness defects associated with Q577R/N/K as well as those that contribute modestly to PIE12-trimer resistance, as W1 and W2 are slightly more resistant than the Q577 mutants alone (Fig. 1 and Table 1).

Using the deep sequencing data, we identified all point mutations, insertions, and deletions within the *env* gene of the PIE12-trimer resistant populations with >10% absolute difference in abundance from the control pool (Table 3 and Supplementary Table 1). We predict that 10% is a high enough threshold to filter out noise due to genetic drift and sequencing errors, but low enough to catch minor variations in the population that could contribute to resistance. Following these guidelines, 25 candidate protein mutations (74 *env* nucleotide positions) were identified for further analysis.

**Table 3:** Amino Acid Changes in HIV-1 Env in Polyclonal Viral Pools with High-Level PIE12-trimer Resistance. List of amino acid mutations (point mutant, insertion or deletion) in either resistance pool that occurs at a rate >10% different than in the control pool. In bold are the mutants that meet the stricter criteria of: 1) high prevalence (≥50%) within both resistant populations but not in the control, 2) not silent (except in the RRE, where a silent mutation could impact RNA structure).

| Mutation | Codon | Prevalence |  |  | Notes |
| --- | --- | --- | --- | --- | --- |
|  |  | Control | W1 | W2 |  |
| T19I | acc → atc | | 44% | | |
| **A48T** | **gca → aca** |  | **72%** | **57%** |  |
| E83K | gaa → aaa | 32% | 20% | 10% | |
| N136D | aat → gat | | 99% | 99% | glycosylation site |
| N136K | aat → aag | 47% | | | |
| R146K | aga → aaa | | 18% | 24% | |
| **Δ161-164** |  |  | **90%** | **90%** | **V1/V2, glycosylation site** |
| F175L | ttc → ctc | | | 67% | V1/V2 |
| V200E | gtc → gaa | | 67% | | |
| V255A | gta → gca | | | 46% | |
| N301K | aac → aaa | 71% | 100% | 99% | glycosylation site |
| S306R | agt → aga | | 64% | 20% | V3 |
| T319A | aca → gca | | 85% | | |
| **Δ396-400** |  |  | **58%** | **59%** | **V4, glycosylation site** |
| N460I | aat → att | | | 17% | near CD4 binding site |
| G464E | ggg → gag | | 82% | | glycosylation site |
| S465P | tcc → ccc | | | 11% | |
| **Q550H** | **cag → cac** |  | **92%** | **98%** | **N-trimer, RRE** |
| **Q577R** | **cag → cgg** |  |  | **99%** | **PIE12 binding site, RRE** |
| **Q577N** | **cag → aac** |  | **97%** |  |  |
| **V583V** | **gtg → gta** |  | **100%** | **100%** | **silent/RRE** |
| A612T | gct → act |  | 13% |  | glycosylation site, RRE |
| H643Y | cac → tac | 99% | 46% | 97% | C-peptide |
| **L663F** | **tta → ttt** |  | **99%** | **99%** |  |
| N674T | aac → acc |  |  | 56% | glycosylation site |
| V693I | gta → ata |  |  | 21% |  |
| **A823V** | **gct → gtt** |  | **98%** | **99%** |  |

To focus our analysis on a more limited set of mutations most likely to contribute to resistance and fitness compensation, we narrowed our list to the those that were not silent (except in the Rev Response Element (RRE), where a silent mutation could impact RNA structure) and with the highest penetrance (>50% abundance in both resistant viral pools but not in the control). Eight mutations meet these criteria (indicated in bold in Table 3). Interestingly, when we compared the clonal Sanger sequence data to these final mutations of interest, all but one are present in every clone. Only A48T lacked 100% penetrance and was found in 3/5 W1 and 3/8 W2 clones.

In addition to identifying the primary resistance mutation (Q577R/N/K) in the PIE12-trimer binding site, this analysis identified three gp120 and four gp41 mutations (Table 3 and Fig. 5). Of the gp120 mutations, one is a point mutation (A48T), and two are in-frame deletions (Δ161-164 in the V1/V2 loop and Δ396-400 in the V4 loop). The V4 loop deletion truncates a tandem duplication of 5 amino acids (FNSTWFNSTW), with only one FNSTW copy linked with PIE12-trimer resistance. This deletion has also been reported in a virus resistant to T-2635, a third-generation C-peptide fusion inhibitor [15]. The four gp41 mutations are all located outside the PIE12 binding site. Two (Q550H and the silent V583V) are located within the RRE, L663F is in the gp41 membrane-proximal external region (MPER), and A823V is in the cytoplasmic domain between LLP-3 and LLP-1.

**Figure 5:**
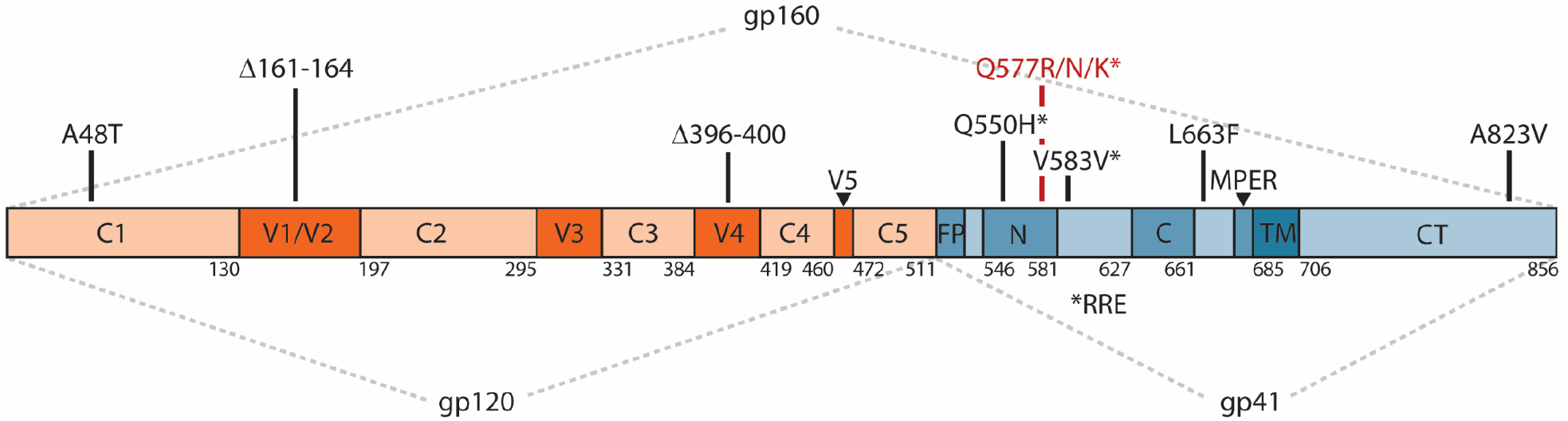
Prominent mutations in gp120 and gp41 observed in PIE12-trimer resistant viruses. The primary structure of the Env precursor, gp160, is depicted, with HXB2 (UniProtKB P04578) numbering. The gp41 N-and C-peptide regions are numbered according to [33]. gp160 is processed into two non-covalently associated subunits, the surface gp120 (orange) and the membrane-spanning gp41 (blue). gp120 comprises five constant domains (C1-C5) and five variable domains (V1-V5). gp41 contains a fusion peptide (FP), an N-peptide region that forms the N-trimer coiled coil (N), a C-peptide region (C) that together with the N-trimer forms a 6-helix bundle in the post-fusion state, a membrane-proximal external region (MPER), a transmembrane domain (TM), and a cytoplasmic tail (CT). The primary resistance mutations (Q577R/N/K) are represented in red, and candidate resistance and fitness compensation mutations are shown in black. Mutations denoted by * are within the Rev Response Element (RRE).

## Discussion

This study identifies Q577R/N/K within the gp41 N-trimer hydrophobic pocket as the primary resistance mechanism used to escape inhibition by the highly potent D-peptide entry inhibitor PIE12-trimer. As predicted by the resistance capacitor hypothesis, the selection for high-level resistance required: 1) extended passaging compared to those against earlier-generation entry inhibitors, 2) a mutation that dramatically reduced inhibitor affinity at the cost of substantially reduced viral fitness, and 3) compensatory mutations to restore viral fitness in the face of such a destabilizing mutation. Despite these obstacles, HIV-1 was ultimately able to resist PIE12-trimer in vitro.

Interestingly, even though the gp41 hydrophobic pocket is highly conserved, in rare cases HIV-1 naturally accommodates alternate residues at position Q577. For example, the uncommon Group O HIV-1 strains (<1% of global HIV-1 infections) primarily contain Q577R. Not surprisingly, we previously observed high-level PIE12-trimer resistance in two HIV-1 Group O isolates [8]. Additionally, within the >50,000 non-Group O sequences from the Los Alamos HIV Sequence Database, ∼1.4% contain Q577R.

Q577R has also been identified in viral resistance screens against other HIV-1 entry inhibitors that target the prehairpin intermediate. Against N-trimer peptides N44, N36, and IZN36, the C-peptide T2635, and the retrocyclin RC-101, Q577R is found in some [15, 34, 35] or all [36, 37] of the resistant viruses and always accompanied by other gp41 mutation(s). By itself, Q577R provided only modest resistance to the N-trimer and C-peptide inhibitors (<3-fold) [15, 37]. Viral escape from a recently identified adnectin, 6200_A08, which similarly targets the gp41 hydrophobic pocket, has also been shown to use Q577R as the primary mechanism [38]. Consistent with our results, many of these studies have demonstrated a fitness defect for Q577R [15, 36–38]. Importantly, Q577R Env (HXB2 strain) has been shown to be properly expressed, processed, and incorporated into pseudovirions, so the fitness defect is likely downstream of virus production [39]. Surprisingly, genetic analysis of the virus from a patient resistant to the FDA-approved peptide fusion inhibitor enfuvirtide also identified the Q577R mutation [40], and in the context of that patient’s pre-treatment *env*, provided 25-fold resistance to enfuvirtide. This discovery is an outlier, however, as the primary resistance mechanism to enfuvirtide involves mutation within gp41 residues 547-556 [11], and the enfuvirtide sensitivity of other studied HIV-1 strains is minimally affected by Q577R [15, 35–37]. Indeed, our data demonstrate only modest differences in C-peptide inhibition of Q577R compared to WT pseudoviruses, including C34, T20 (enfuvirtide) and T1249 (Supplementary Table 2), and our PIE12-trimer resistant pools retain sensitivity to a related peptide, C37 (Table 1).

The Q577 mutation in the critical N-trimer region could conceivably influence multiple stages of the entry process, leading to its effect on viral fitness. Using multiple methods, Weiss, et al. have investigated the effect of Q577R on Env structure and function. They have shown Q577R leads to: 1) a more stable 6-helix bundle structure, 2) greater neutralization sensitivity to CD4 mimetics, and 3) decreased susceptibility to CD4-induced gp41 conformational changes that lead to viral inactivation [34, 35, 37, 41]. This wide influence of Q577 mutation on Env function supports the concept that both gp120 and gp41 modifications would be needed to restore viral fitness, as is seen in our PIE12-trimer resistant pools. However, there is little overlap between the Env modifications seen in our pools and those in naturally occurring strains with Q577R and in Q577R virus resistant to other entry inhibitors. As shown in an alignment of the ∼750 Group M primary isolate Q577R Envs (out of >50,000 Group M sequences from the Los Alamos HIV Sequence Database) with our PIE12-trimer resistant sequences, none of the candidate PIE12-trimer-resistant compensatory mutations are prevalent in the primary isolates (Supplementary Table 3). Additionally, we did not identify any overlap of our specific compensatory mutations with those found in the other entry-inhibitor resistance screens with Q577R virus [15, 35–37].

Although none of our specific compensatory mutations are found in Q577R viruses resistant to other entry inhibitors, it seems likely that similar compensation mechanisms are used. In previous entry inhibitor resistance studies that include Q577R and for which gp120 sequence is available [15, 34–36, 41], compensatory mutations are found in CD4 binding residues (E426K), the V1/V2 loop (D167N), and in the V3 loop (R298K, S306G, I309M, T319I, and E322D), indicating that changes to gp120-based fusion determinants, such as CD4 and co-receptor binding, help restore gp41-based fitness defects. Our V1/V2 and V4 loop deletions may similarly affect Env function. Additionally, several V3 loop mutants that are similar to those identified in other resistance selections were found in some of our sequences (S306R, T319A).

The region of the HIV genome that encodes the highly conserved hydrophobic pocket (the target of PIE12) also encodes part of the Rev-Response Element (RRE), whose specific RNA structure is required for proper viral replication. Previous work showed that silent mutations at Q550 and Q577, as well as the double mutant, are sufficient to induce structural changes in the RRE [42]. Interestingly, Q550H and Q577R/N/K are observed in nearly all PIE12-trimer resistant sequences. We postulate that the primary resistance mutation Q577, which is located at the base of stem-loop V [43], forces a structural rearrangement of this conserved RRE structure and contributes to the fitness defect. The presence of additional mutations within the RRE, namely Q550H and a silent mutation at V583(GTG→GTA), suggest these may play an important compensatory role in restructuring the RRE. Q550 mutations, however, are not observed in any of the 689 Q577R-containing primary isolates (Supplementary Table 3). Residue 583 is variable in the Q577R-containing primary isolates, although only hydrophobic residues (valine, isoleucine, leucine, and methionine) are observed.

Compensatory mutations to changes in the RRE could also be found in Rev, which binds the RRE to promote the export of mRNA from the nucleus. Although our sequence analysis focused on mutations in *env*, 80% of the Rev sequence (exon 2) is encoded within this same genomic region. Three Rev mutations (*env* positions 2354, 2375 and 2435, leading to N86Y, G93W, and G113R in Rev while silent in Env) surfaced in our initial analysis (Supplementary Table 1), but each is present in only one resistant pool, none are fully penetrant, and all are located in the predicted disordered nuclear export sequence [44]. Sequencing of *rev* exon 1 and functional studies of mutant Rev/RRE will be required to see if there are any meaningful compensatory changes in Rev.

We have also sought to improve the overall potency and breadth of PIE12-trimer by conjugating it to cholesterol (chol-PIE12-trimer) to concentrate it at cell membrane lipid rafts, the site of viral entry. This modification increases its association rate for the prehairpin intermediate and improves potency ∼100-fold [10]. Additionally, cholesterol conjugation prolongs the in vivo half-life of the inhibitor, making it suitable for extended-release delivery [45]. With its improved potency, chol-PIE12-trimer is better able to combat PIE12-trimer resistant virus and inhibits pseudovirions with the Q577R mutation with a low nanomolar IC_50_ [10]. However, our preliminary viral passaging studies in the presence of chol-PIE12-trimer suggest high-level resistant virus ultimately emerges. Like for the PIE12-trimer resistant viruses, the pathway to resistance against chol-PIE12-trimer includes mutation of Q577, and no other pocket mutations are observed. We are currently studying the additional mutations acquired during chol-PIE12-trimer passaging to understand the mechanism of resistance against this potential clinical candidate.

Ultimately, we hope to use the data described here to develop next-generation D-peptide inhibitors that anticipate this route of resistance to create an even higher barrier to resistance. Efforts are underway to discover novel D-peptide entry inhibitors that maintain high affinity to both the native and Q577R pockets.

## Conclusion

Although our highly potent, broad D-peptide entry inhibitor, PIE12-trimer, exhibits a high barrier to resistance, with extended passaging (>1 year) we obtained HIV-1 resistant to PIE12-trimer. The primary resistance mutation, Q577R/N/K, is found in the conserved PIE12-trimer binding pocket on the gp41 N-trimer. Q577R disrupts two hydrogen bonds between PIE12-trimer and the inhibitor-binding pocket and substantially decreases binding affinity. Q577 mutation confers a significant fitness defect as seen by delayed viral replication. The highly resistant polyclonal viral pools exhibit rescued fitness due to compensatory mutations found in gp120 and gp41. Using deep sequencing analysis, we identified seven potential compensatory mutations which will be the focus of future mechanistic studies. Possible compensatory mechanisms include modifications to viral entry steps, such as receptor and co-receptor binding, fusion kinetics, Env folding, and interactions between Rev and the RRE.

## Declarations

### Ethics Approval and Consent to Participate

Not applicable

### Consent for Publication

Not applicable.

### Availability of Data and Materials

Deep-sequence data from the polyclonal viral pools and Perl scripts used to process them available upon request. Coordinates for the PIE12/IQN17-Q577R complex structure are available at the protein data bank (PDB code: 6PSA).

### Author’s contributions

ARS and MTW spearheaded the project including experimental plans, data analysis, and manuscript preparation. MTW conducted the biophysical characterization, crystallography studies, viral genomics, and bioinformatics analysis. ARS conducted the viral characterization, inhibition and growth studies. AES conducted the viral passaging studies. FGW assisted with x-ray crystallography data collection and analysis. JNF contributed to the virology studies. DME contributed significantly to project design and manuscript preparation. CPH supervised x-ray crystallography studies. MJR supervised the viral passaging studies. MSK provided guidance for the overall project.

## Acknowledgements

The authors would like to gratefully acknowledge Ethan B. Howell for his assistance with binding affinity measurements, Siyu Chen for molecular biology, and Brian Dalley (High-Throughput Genomics and Bioinformatic Analysis, Huntsman Cancer Institute, University of Utah) for deep-sequencing advice and services (supported by the National Cancer Institute of the National Institutes of Health under Award Number P30CA042014). Use of the Stanford Synchrotron Radiation Lightsource (SSRL), SLAC National Accelerator Laboratory, is supported by the U.S. Department of Energy, Office of Science, Office of Basic Energy Sciences under Contract No. DE-AC02-76SF00515. The SSRL Structural Molecular Biology Program is supported by the DOE Office of Biological and Environmental Research, and by the National Institutes of Health, National Institute of General Medical Sciences (including P41GM103393). The contents of this publication are solely the responsibility of the authors and do not necessarily represent the official views of NIGMS or NIH.

## Authors’ Information

The current affiliations of authors are as follows: ARS (Utah Department of Health, Salt Lake City, UT), MTW (Group Leader, Molecular Sciences, AbSci, LLC, Vancouver, WA), JNF (BioFire Diagnostics, Salt Lake City).

