## Supplementary Material for "Characterization of resistance to a potent D-peptide HIV entry inhibitor"

Supplementary Figure 1: SPR sensorgrams for PIE12 monomer binding to IZN36 WT (top panel) or Q577R (bottom panel), processed in Scrubber2 (BioLogic Software) and used for the equilibrium fit shown in Figure 2. Six 3-fold dilutions of PIE12 monomer (617 nM to 2.54 nM) were flowed over the WT surface, and ten 3-fold dilutions (50 uM to 2.54 nM) were flowed over the Q577R surface. The calculated $K_D$'s are 0.031 μM for WT and 2.0 μM for Q577R.

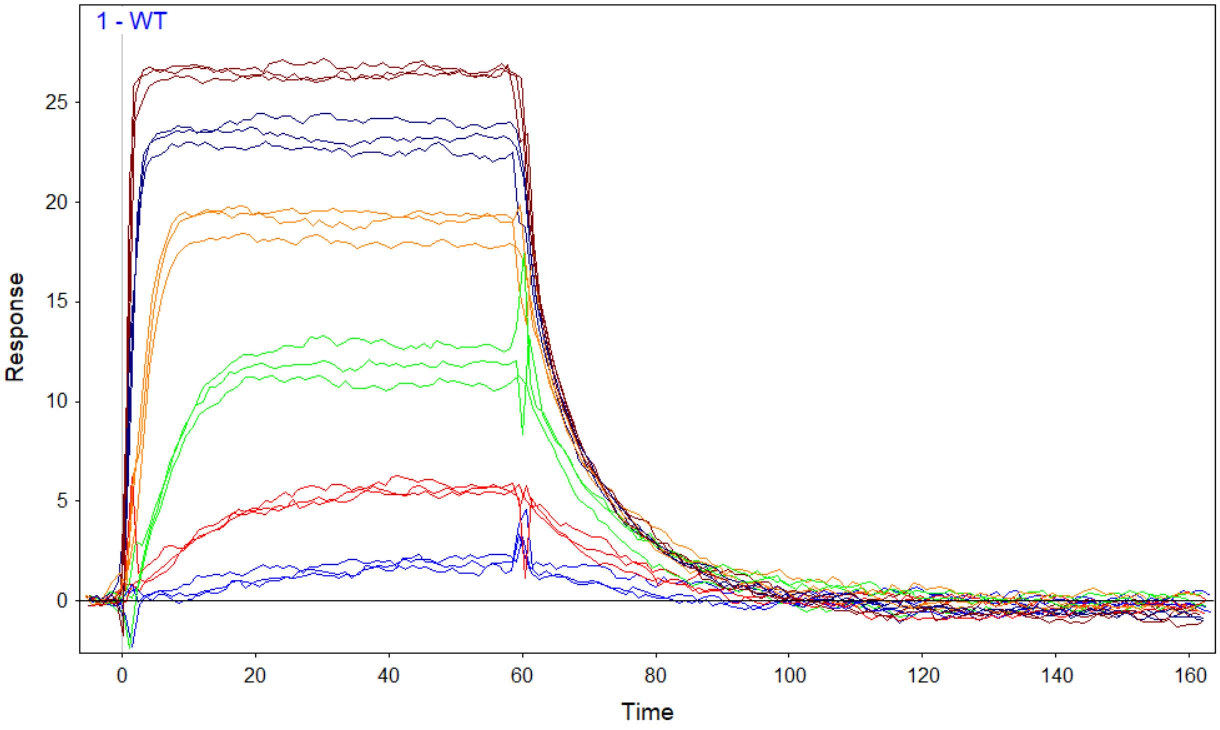


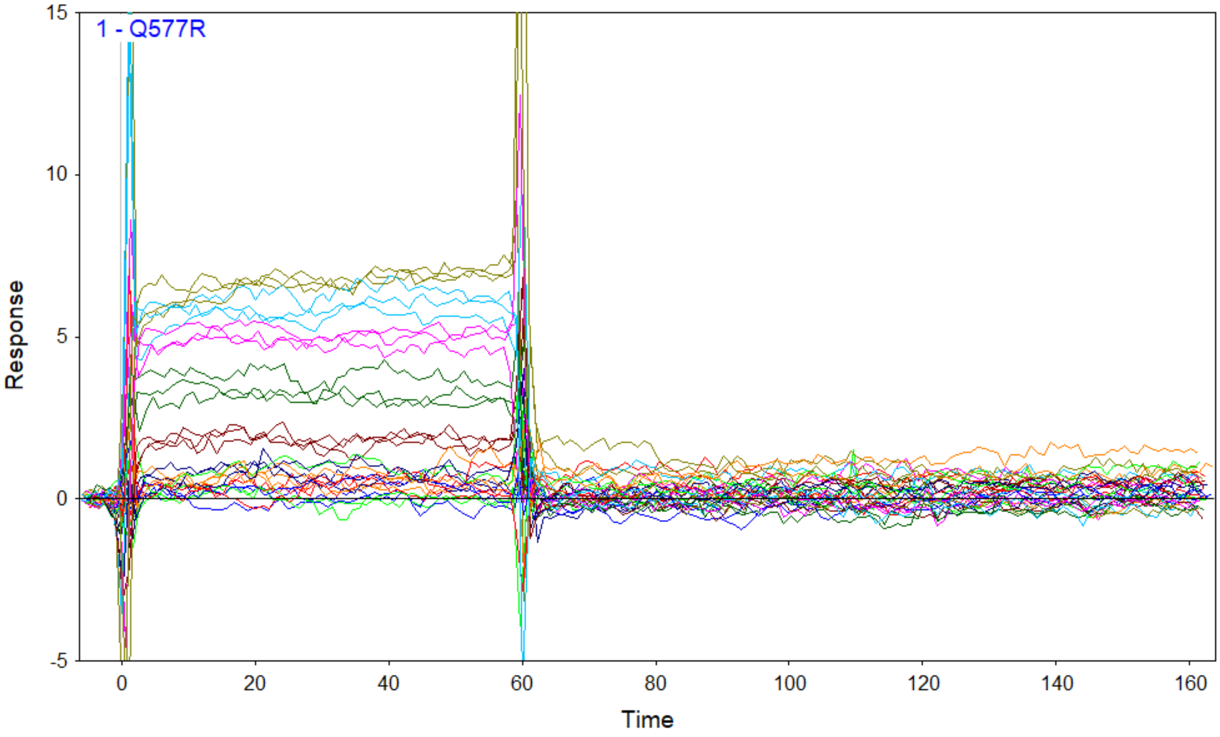

Supplementary Table 1: Mutations observed in PIE12-trimer resistant viral pools. *Env* gene position (pNL4-3 numbering), reference base, and all base calls for control and PIE12-trimer resistance viral populations with a mutation >10% of the population (with percentage of the base calls for each position colored on a spectrum from low (light pink) to high (red)) are indicated. Additionally, the positions are annotated by their location in Env according to the following categories: known or putative glycosylation site (green), gp120 variable loops (gray), chemokine receptor (co-receptor) binding site (blue), rev-response element (dark gray), gp41 N-trimer (orange), gp41 C-peptide region (light blue), or the Rev coding region included within the *env* gene (yellow). Eighty-one positions were identified. Of these, 74 conform to our analysis criteria (found in the resistant pool(s) and >10% different frequency than in the control pool).

| | | | Control | | | | | | 9754X8 320nM PIE12-trimer W1 | | | | | | 9754X9 320nM PIE12-trimer W2 | | | | | |
|---|---|---|---|---|---|---|---|---|---|---|---|---|---|---|---|---|---|---|---|---|
| POS | REF | annotation | A | C | G | T | IN | DEL | A | C | G | T | IN | DEL | A | C | G | T | IN | DEL |
| 82 | c | | | | | | | | 0.0 | 55.6 | 0.0 | 44.3 | 0.0 | 0.0 | | | | | | |
| 140 | c | | 11.5 | 87.7 | 0.0 | 0.6 | 0.0 | 0.0 | | | | | | | | | | | | |
| 168 | g | | | | | | | | 72.0 | 0.1 | 27.7 | 0.0 | 0.0 | 0.0 | 56.9 | 0.1 | 42.9 | 0.0 | 0.0 | 0.0 |
| 273 | g | | 31.7 | 0.0 | 68.2 | 0.0 | 0.0 | 0.0 | 20.5 | 0.0 | 79.4 | 0.0 | 0.0 | 0.0 | 10.1 | 0.0 | 89.8 | 0.0 | 0.0 | 0.0 |
| 280 | t | | 0.1 | 0.2 | 0.1 | 76.6 | 23.0 | 0.0 | 0.1 | 0.1 | 0.1 | 84.1 | 15.6 | 0.0 | | | | | | |
| 285 | g | | 0.6 | 0.1 | 76.8 | 0.1 | 22.3 | 0.1 | 0.5 | 0.1 | 84.2 | 0.1 | 15.0 | 0.1 | | | | | | |
| 432 | a | glyco | | | | | | | 0.6 | 0.0 | 99.2 | 0.0 | 0.0 | 0.1 | 0.8 | 0.0 | 98.9 | 0.1 | 0.0 | 0.1 |
| 463 | g | V1/V2 | | | | | | | 17.7 | 0.0 | 82.2 | 0.0 | 0.0 | 0.0 | 24.3 | 0.9 | 74.7 | 0.0 | 0.0 | 0.0 |
| 476 | g | V1/V2 | | | | | | | 14.5 | 0.0 | 85.3 | 0.0 | 0.0 | 0.0 | | | | | | |
| 508 | t | V1/V2 | | | | | | | 0.0 | 0.0 | 0.0 | 53.0 | 0.0 | 46.9 | 0.0 | 0.0 | 0.0 | 53.0 | 0.1 | 46.8 |
| 509 | c | V1/V2 | | | | | | | 9.5 | 0.1 | 0.0 | 0.0 | 0.0 | 90.3 | 9.7 | 0.1 | 0.0 | 0.0 | 0.0 | 90.1 |
| 510 | a | V1/V2 | | | | | | | 7.7 | 0.0 | 0.0 | 0.0 | 0.0 | 92.3 | 7.8 | 0.0 | 0.0 | 0.0 | 0.0 | 92.1 |
| 511 | g | V1/V2 | | | | | | | 0.0 | 0.0 | 5.3 | 0.0 | 0.0 | 94.6 | 0.0 | 0.0 | 5.5 | 0.0 | 0.0 | 94.4 |
| 512 | c | V1/V2 | 99.6 | 0.0 | 0.1 | 0.0 | 0.0 | 0.1 | 2.8 | 0.0 | 0.0 | 0.0 | 0.0 | 97.1 | 2.9 | 0.0 | 0.0 | 0.0 | 0.0 | 97.0 |
| 513 | a | V1/V2 | | | | | | | 0.1 | 0.0 | 0.2 | 0.0 | 0.0 | 99.6 | 0.1 | 0.0 | 0.2 | 0.0 | 0.0 | 99.6 |
| 514 | c | V1/V2 | | | | | | | 0.2 | 0.1 | 0.0 | 0.0 | 0.0 | 99.7 | 0.2 | 0.1 | 0.0 | 0.0 | 0.0 | 99.6 |
| 515 | a | V1/V2 | | | | | | | 0.1 | 0.0 | 0.0 | 0.1 | 0.0 | 99.7 | 0.2 | 0.0 | 0.0 | 0.1 | 0.0 | 99.6 |
| 516 | a | V1/V2 | | | | | | | 3.0 | 0.0 | 0.0 | 0.0 | 0.0 | 97.0 | 3.1 | 0.0 | 0.0 | 0.0 | 0.0 | 96.8 |
| 517 | g | V1/V2 | | | | | | | 4.9 | 0.0 | 0.1 | 0.0 | 0.0 | 94.9 | 4.9 | 0.0 | 0.2 | 0.0 | 0.0 | 94.9 |
| 518 | c | V1/V2 | | | | | | | 0.0 | 2.7 | 0.1 | 7.0 | 0.0 | 90.1 | 0.0 | 3.3 | 0.1 | 7.0 | 0.0 | 89.6 |
| 519 | a | V1/V2 | | | | | | | 11.8 | 0.0 | 0.1 | 0.0 | 2.5 | 85.6 | 12.4 | 0.0 | 0.0 | 0.0 | 2.6 | 84.9 |
| 520 | t | V1/V2 | | | | | | | 0.0 | 0.0 | 0.0 | 14.1 | 0.0 | 85.8 | 0.0 | 0.0 | 0.0 | 14.8 | 0.0 | 85.1 |
| 549 | t | V1/V2 | | | | | | | | | | | | | 0.0 | 66.6 | 0.0 | 33.2 | 0.0 | 0.1 |
| 619 | t | | 99.4 | 0.1 | 0.1 | 0.2 | 0.0 | 0.1 | 65.8 | 0.1 | 0.0 | 34.0 | 0.0 | 0.0 | | | | | | |
| 620 | c | | | | | | | | 65.5 | 34.2 | 0.0 | 0.2 | 0.0 | 0.0 | | | | | | |
| 683 | t | | | | | | | | 0.0 | 32.7 | 0.0 | 67.1 | 0.0 | 0.0 | | | | | | |
| 784 | t | | | | | | | | | | | | | | 0.0 | 46.4 | 0.1 | 53.4 | 0.0 | 0.0 |
| 884 | g | | | | | | | | | | | | | | 11.6 | 0.0 | 88.3 | 0.0 | 0.0 | 0.0 |
| 923 | c | glyco | 70.7 | 27.2 | 1.7 | 0.2 | 0.0 | 0.1 | 99.6 | 0.1 | 0.1 | 0.0 | 0.0 | 0.1 | 99.4 | 0.1 | 0.1 | 0.2 | 0.0 | 0.1 |
| 938 | t | V3 | | | | | | | 64.2 | 0.1 | 0.1 | 35.5 | 0.0 | 0.0 | 20.3 | 0.1 | 0.2 | 79.3 | 0.0 | 0.1 |
| 972 | g | V3 | | | | | | | | | | | | | 39.8 | 0.0 | 60.1 | 0.0 | 0.0 | 0.0 |
| 975 | a | V3 | | | | | | | 15.2 | 0.0 | 84.5 | 0.0 | 0.1 | 0.1 | | | | | | |
| 1043 | t | | | | | | | | | | | | | | 0.6 | 19.5 | 0.0 | 79.7 | 0.0 | 0.0 |
| 1179 | t | V4 | | | | | | | | | | | | | 17.4 | 0.0 | 0.0 | 82.5 | 0.0 | 0.0 |
| 1205 | g | V4 | | | | | | | 0.2 | 0.0 | 58.0 | 0.0 | 2.4 | 39.3 | 0.1 | 0.0 | 57.6 | 0.0 | 2.4 | 39.8 |
| 1206 | t | V4 | | | | | | | 7.3 | 0.0 | 0.1 | 24.4 | 0.0 | 68.1 | 6.9 | 0.0 | 0.1 | 22.7 | 0.0 | 70.1 |
| 1207 | t | V4 | | | | | | | 0.1 | 0.0 | 4.6 | 25.3 | 0.0 | 69.9 | 0.0 | 0.0 | 4.3 | 23.6 | 0.0 | 71.9 |
| 1208 | t | V4 | | | | | | | 0.0 | 0.0 | 0.2 | 31.5 | 0.0 | 68.2 | 0.0 | 0.1 | 0.1 | 29.6 | 0.0 | 70.1 |
| 1209 | a | glyco | | | | | | | 33.1 | 0.0 | 0.1 | 0.1 | 0.0 | 66.7 | 31.8 | 0.0 | 0.0 | 0.0 | 0.0 | 68.1 |
| 1210 | a | glyco | | | | | | | 33.1 | 0.3 | 0.0 | 0.0 | 0.0 | 66.5 | 32.2 | 0.3 | 0.0 | 0.0 | 0.0 | 67.4 |
| 1211 | t | glyco | | | | | | | 0.0 | 0.3 | 0.0 | 34.8 | 0.0 | 64.8 | 0.3 | 0.1 | 0.0 | 33.6 | 0.0 | 65.9 |
| 1212 | a | V4 | | | | | | | 41.7 | 0.0 | 0.2 | 0.1 | 0.0 | 57.9 | 40.7 | 0.0 | 0.1 | 0.0 | 0.0 | 59.0 |
| 1213 | g | V4 | | | | | | | 0.2 | 0.1 | 43.1 | 0.1 | 0.0 | 56.4 | 0.1 | 0.1 | 42.1 | 0.0 | 0.0 | 57.6 |
| 1214 | t | V4 | | | | | | | 0.1 | 0.2 | 0.1 | 44.8 | 0.0 | 54.7 | 0.1 | 0.1 | 0.1 | 43.8 | 0.0 | 55.9 |
| 1215 | a | V4 | | | | | | | 46.6 | 0.0 | 0.2 | 0.0 | 0.0 | 53.1 | 45.4 | 0.0 | 0.1 | 0.0 | 0.0 | 54.3 |
| 1216 | c | V4 | | | | | | | 0.1 | 48.1 | 0.1 | 0.1 | 0.0 | 51.6 | 0.0 | 47.0 | 0.1 | 0.2 | 0.0 | 52.7 |
| 1217 | t | V4 | | | | | | | 0.1 | 0.1 | 0.1 | 48.0 | 0.0 | 51.5 | 0.1 | 0.2 | 0.1 | 46.8 | 0.0 | 52.8 |
| 1218 | t | V4 | | | | | | | 0.1 | 0.0 | 0.2 | 48.2 | 0.0 | 51.5 | 0.1 | 0.0 | 0.2 | 46.9 | 0.0 | 52.8 |
| 1219 | g | V4 | | | | | | | 0.2 | 0.0 | 49.5 | 0.0 | 0.0 | 50.2 | 0.1 | 0.0 | 48.3 | 0.0 | 0.0 | 51.4 |
| 1220 | g | V4 | | | | | | | 0.1 | 0.1 | 50.3 | 0.1 | 0.0 | 49.4 | 0.1 | 0.0 | 49.1 | 0.0 | 0.0 | 50.7 |
| 1351 | g | CKR | | | | | | | 12.6 | 0.0 | 87.3 | 0.0 | 0.0 | 0.0 | | | | | | |
| 1399 | a | | | | | | | | | | | | | | 83.3 | 0.1 | 0.0 | 16.5 | 0.0 | 0.0 |
| 1411 | g | | | | | | | | 81.7 | 0.0 | 18.1 | 0.0 | 0.0 | 0.0 | | | | | | |
| 1413 | t | | | | | | | | | | | | | | 0.0 | 11.1 | 0.1 | 88.7 | 0.0 | 0.0 |
| 1617 | c | RRE | 0.1 | 4.4 | 95.4 | 0.0 | 0.0 | 0.0 | 0.1 | 4.6 | 95.2 | 0.0 | 0.0 | 0.0 | 0.1 | 4.4 | 95.4 | 0.0 | 0.0 | 0.1 |
| 1618 | g | RRE | 0.0 | 93.9 | 4.2 | 0.1 | 1.7 | 0.0 | 0.0 | 93.9 | 4.1 | 0.1 | 1.7 | 0.0 | 0.0 | 93.9 | 4.1 | 0.1 | 1.7 | 0.1 |
| 1660 | a | N-pep | 0.2 | 0.0 | 99.7 | 0.0 | 0.0 | 0.0 | 0.1 | 0.0 | 99.8 | 0.0 | 0.0 | 0.0 | 0.1 | 0.0 | 99.7 | 0.0 | 0.0 | 0.0 |
| 1670 | g | N-pep | | | | | | | 0.0 | 92.1 | 0.1 | 7.7 | 0.0 | 0.0 | 0.0 | 98.0 | 0.1 | 1.8 | 0.0 | 0.0 |
| 1676 | g | N-pep | 24.1 | 0.0 | 75.8 | 0.0 | 0.0 | 0.0 | 14.4 | 0.0 | 85.5 | 0.0 | 0.0 | 0.0 | | | | | | |
| 1749 | c | N-pep | | | | | | | 97.3 | 2.5 | 0.2 | 0.0 | 0.0 | 0.0 | | | | | | |
| 1750 | a | N-pep | | | | | | | | | | | | | 0.4 | 0.0 | 99.5 | 0.0 | 0.0 | 0.0 |
| 1751 | g | N-pep | | | | | | | 0.1 | 97.4 | 1.3 | 1.0 | 0.0 | 0.0 | | | | | | |
| 1769 | g | RRE | | | | | | | 99.8 | 0.0 | 0.1 | 0.0 | 0.0 | 0.0 | 99.7 | 0.0 | 0.1 | 0.0 | 0.0 | 0.0 |
| 1854 | g | RRE | | | | | | | 13.1 | 0.0 | 86.8 | 0.0 | 0.0 | 0.0 | | | | | | |
| 1901 | c | | | | | | | | | | | | | | 11.4 | 87.5 | 0.0 | 1.0 | 0.0 | 0.0 |
| 1947 | c | C-pep | 0.0 | 0.8 | 0.0 | 99.1 | 0.0 | 0.0 | 0.0 | 53.6 | 0.0 | 46.3 | 0.0 | 0.0 | 0.0 | 2.9 | 0.0 | 96.9 | 0.0 | 0.0 |
| 2009 | a | | | | | | | | 0.5 | 0.1 | 0.0 | 99.1 | 0.0 | 0.2 | 0.6 | 0.1 | 0.0 | 99.1 | 0.0 | 0.1 |
| 2041 | a | putative glyco | | | | | | | | | | | | | 43.3 | 56.5 | 0.1 | 0.0 | 0.0 | 0.0 |
| 2097 | g | | | | | | | | | | | | | | 21.3 | 0.0 | 78.6 | 0.0 | 0.0 | 0.0 |
| 2135 | g | | | | | | | | | | | | | | 20.1 | 0.0 | 79.7 | 0.0 | 0.0 | 0.0 |
| 2354 | a | rev | | | | | | | | | | | | | 34.5 | 0.1 | 0.0 | 65.3 | 0.0 | 0.0 |
| 2375 | g | rev | | | | | | | | | | | | | 0.3 | 0.1 | 42.0 | 57.5 | 0.0 | 0.0 |
| 2435 | g | rev | | | | | | | 20.5 | 0.0 | 79.3 | 0.1 | 0.0 | 0.0 | | | | | | |
| 2450 | t | | | | | | | | | | | | | | 41.7 | 0.1 | 0.2 | 57.9 | 0.0 | 0.1 |
| 2452 | c | | | | | | | | | | | | | | 0.0 | 89.4 | 0.1 | 10.4 | 0.0 | 0.0 |
| 2488 | c | | | | | | | | 0.2 | 1.4 | 0.1 | 98.2 | 0.0 | 0.0 | 0.1 | 0.5 | 0.1 | 99.2 | 0.0 | 0.0 |
| 2499 | g | | | | | | | | 16.0 | 0.3 | 83.6 | 0.1 | 0.0 | 0.0 | | | | | | |
| 2505 | g | | | | | | | | | | | | | | 18.5 | 0.0 | 81.3 | 0.0 | 0.0 | 0.1 |
| 2573 | g | | | | | | | | | | | | | | 25.0 | 0.0 | 74.7 | 0.1 | 0.0 | 0.1 |
| 2579 | g | | | | | | | | 25.2 | 0.0 | 74.5 | 0.0 | 0.0 | 0.2 | 34.3 | 0.0 | 65.5 | 0.0 | 0.0 | 0.1 |
| 2599 | g | | | | | | | | 14.2 | 0.0 | 85.7 | 0.0 | 0.0 | 0.0 | | | | | | |

Supplementary Table 2: Effect of Q577R on C-peptide Inhibitors

Single-cycle viral infectivity assays in which HIV-1 HXB2 Env (WT and Q577R) pseudotyped HIV-1 with a luciferase reporter was used to infect HOS-LES cells in the absence or presence of six 5-fold dilutions of the indicated C-peptide (in quadruplicate). The data are the average of two experiments with the standard deviation in parentheses.

| | IC50 (nM) | | |
|---|---|---|---|
| | **WT** | **Q577R** | **Fold Difference** |
| **C34** | 0.355 (0.177) | 1.58 (0.30) | 4.8 |
| **T20** | 1.62 (0.23) | 5.33 (0.74) | 3.4 |
| **T1249** | 0.152 (0.064) | 0.18 (0.05) | 1.2 |

Supplementary Table 3: Prevalence of PIE12-trimer resistant candidate compensatory amino acid mutations in Group M primary isolates containing Q577R.

| WT Position | PIE12-trimer resistant Mutation | Prevalence in Q577R containing Primary Isolates | Other mutations observed at this position | Comments |
|---|---|---|---|---|
| A48 | A48**T** | 1/751 | None | • Alanine is conserved in 750/751 (99.9%) sequences at this position |
| 161-164 amino acids: ISTS | Δ161-164 | 0/751 | | • Loss of a glycosylation site with this deletion<br>• 724/751 sequences have the intact glycosylation site |
| 396-400 amino acids: FNSTW | Δ396-400 | 0/751 | | • Loss of a glycosylation site with this deletion<br>• This region is highly variable<br>• 16/751 lack at least one of the V4 glycosylation sites in this region |
| Q550 | Q550**H** | 0/751 | | • The WT Q550 is conserved in all 751 sequences. Twenty-two have the cag codon; 729 have caa. |
| V583 Codon: GTG | V583 Codon: GT**A** | 28/751 | V583I, V583L and V583M (168/751) | • 583/751 sequences have V583. Of these, 28 have the gta codon.<br>• Only hydrophobic residues are seen at this position in this viral pool<br>• All variants at this position (Val, Ile, Leu and Met) have a T as the second position in the codon<br>• In the RRE structure, that T(U) is in a G-U pair at the end of a stem-loop—suggests this T(U) is important |
| L663 | L663**F** | 1/751 | L663W (2/751) | • 748/751 have WT L663 |

| | | | | |
|---|---|---|---|---|
| A823 | A823**V** | 0/751 | A823G (134/751) | • 617/751 have WT A823 |
